# Data-Driven Analysis of a Mechanistic Model of CAR T Cell Signaling Predicts Effects of Cell-to-cell Heterogeneity

**DOI:** 10.1101/808626

**Authors:** Colin G. Cess, Stacey D. Finley

## Abstract

Due to the variability of protein expression, cells of the same population can exhibit different responses to stimuli. It is important to understand this heterogeneity at the individual level, as population averages mask these underlying differences. Using computational modeling, we can interrogate a system much more precisely than by using experiments alone, in order to learn how the expression of each protein affects a biological system. Here, we examine a mechanistic model of CAR T cell signaling, which connects receptor-antigen binding to MAPK activation, to determine intracellular modulations that can increase cellular response. CAR T cell cancer therapy involves removing a patient’s T cells, modifying them to express engineered receptors that can bind to tumor-associated antigens to promote cell killing, and then injecting the cells back into the patient. This population of cells, like all cell populations, would have heterogeneous protein expression, which could affect the efficacy of treatment. Thus, it is important to examine the effects of cell-to-cell heterogeneity. We first generated a dataset of simulated cell responses via Monte Carlo simulations of the mechanistic model, where the initial protein concentrations were randomly sampled. We analyzed the dataset using partial least-squares modeling to determine the relationships between protein expression and ERK phosphorylation, the output of the mechanistic model. Using this data-driven analysis, we found that only the expressions of proteins relating directly to the receptor and the MAPK cascade, the beginning and end of the network, respectively, are relevant to the cells’ response. We also found, surprisingly, that increasing the amount of receptor present can actually inhibit the cell’s ability to respond due to increasing the strength of negative feedback from phosphatases. Overall, we have combined data-driven and mechanistic modeling to generate detailed insight into CAR T cell signaling.

## 1 INTRODUCTION

Even among cells of the same type, phenotypic differences can arise due to variations in protein abundance, which are caused by the stochastic nature of gene expression (Fraser and Kaern, 2009; Mantzaris, 2007; Niepel et al., 2009). While the response of a signaling network may be robust to the variations in the expressions of some proteins, the expressions of other proteins may be highly influential, causing variances that compound into significant differences at the cellular level (Altschuler and Wu, 2010). Although methods such as flow cytometry can measure protein expression in individual cells, it is still difficult to examine experimentally all of the proteins in a network and how they relate to network response. Using computational modeling, it is possible to determine how variations in protein expression affect phenotypic outcome by precisely controlling protein amounts and simulating their effects.

Computational mechanistic models comprised of ordinary differential equations can be analyzed using various approaches. Some methods of analyzing these models, such as sensitivity analysis, require a large computational cost to be performed in all dimensions. The computational resources required to analyze such models can be prohibitive, especially as model size and complexity increases. As an alternative, data-driven methods can be used as a way of analyzing a model in all possible dimensions at once. While data-driven methods are unable to model actual biological interactions, they are able to generalize the relationships between model inputs and outputs, providing information on how each input acts across all dimensions. For example, Hua et al. used a decision tree to analyze the effects of differences in protein expression in a model of Fas-mediated caspase-3 activation. This allowed the authors to see how individual proteins worked together to influence the response of the system (Hua et al., 2006). However, a major drawback to this approach is that as the number of proteins in the network increases, the decision tree must expand as well, containing many more nodes and branches until it becomes very difficult to analyze. We propose here to use partial least-squares (PLS) as an alternative way to analyze a mechanistic model.

PLS provides information on how the inputs of a system, in this case the initial protein expressions, relate to the system’s outputs. Although PLS does not give information about specific relationships between proteins, it does tell how the expression of each protein generally affects the response, providing information on the population as a whole. It is also easy to analyze, with multiple quantitative metrics providing information on the influence of the inputs.

In this study, we apply PLS to a mechanistic model of chimeric antigen receptor (CAR) T cell signaling. The model connects receptor-antigen binding to the MAPK cascade, resulting in the phosphorylation of ERK, which is one characteristic of T cell activation. CAR T cells are a type of cell-based immunotherapy in which T cells that have been taken from a cancer patient are modified to express the CAR on the cell surface, such that the cells can directly bind to the tumor-associated antigen that the CAR recognizes. Once put back into the patient, these engineered T cells can be directly activated by tumor cells, allowing the CAR T cells to kill the diseased cells without targeting other cells in the body (Androulla and Lefkothea, 2018; Sharpe and Mount, 2015). However, this therapeutic approach has some limitations; for example, there are many instances of no response from the patient (Makita et al., 2017). Many different types of CARs have been developed to try to increase tumor killing (Androulla and Lefkothea, 2018; Cho et al., 2018). Most of those modifications focus on the CAR itself (i.e., creating a new receptor with different signaling domains) and not the downstream signaling network. Using PLS, we focus on intracellular variations of proteins involved in a signaling network that influences how a CAR T cell responds to stimulation. We use Monte Carlo simulations to generate a dataset with protein expressions as the inputs and ability to respond to stimulation, based on ERK phosphorylation, as the output. We find that only a small subset of proteins in the network, those relating to the receptor and to the MAPK cascade, have expressions that significantly influence the network’s response.

## 2 METHODS

### Mechanistic Model of CAR T Cell Signaling

This study employed the use of a mechanistic model of CAR T cell signaling that was previously developed (Rohrs et al., 2019). This model was constructed using a modular approach with smaller models that account for lymphocyte-specific protein tyrosine kinase (LCK) regulation, CAR phosphorylation, LAT signalosome formation, CD45 phosphatase activity, mitogen-activated protein kinase (MAPK) activation, and feedback from the phosphatase SHP1. The model structure is shown in Figure 1. Using this model, we are able to simulate T cell signaling initiated by antigen binding to the extracellular domain of the CAR and culminating with activation of the MAPK pathway to produce doubly phosphorylated ERK (ppERK). Here, we consider the concentration of ppERK as the primary model prediction, and it is the focus of all model simulations.

**Figure 1:**
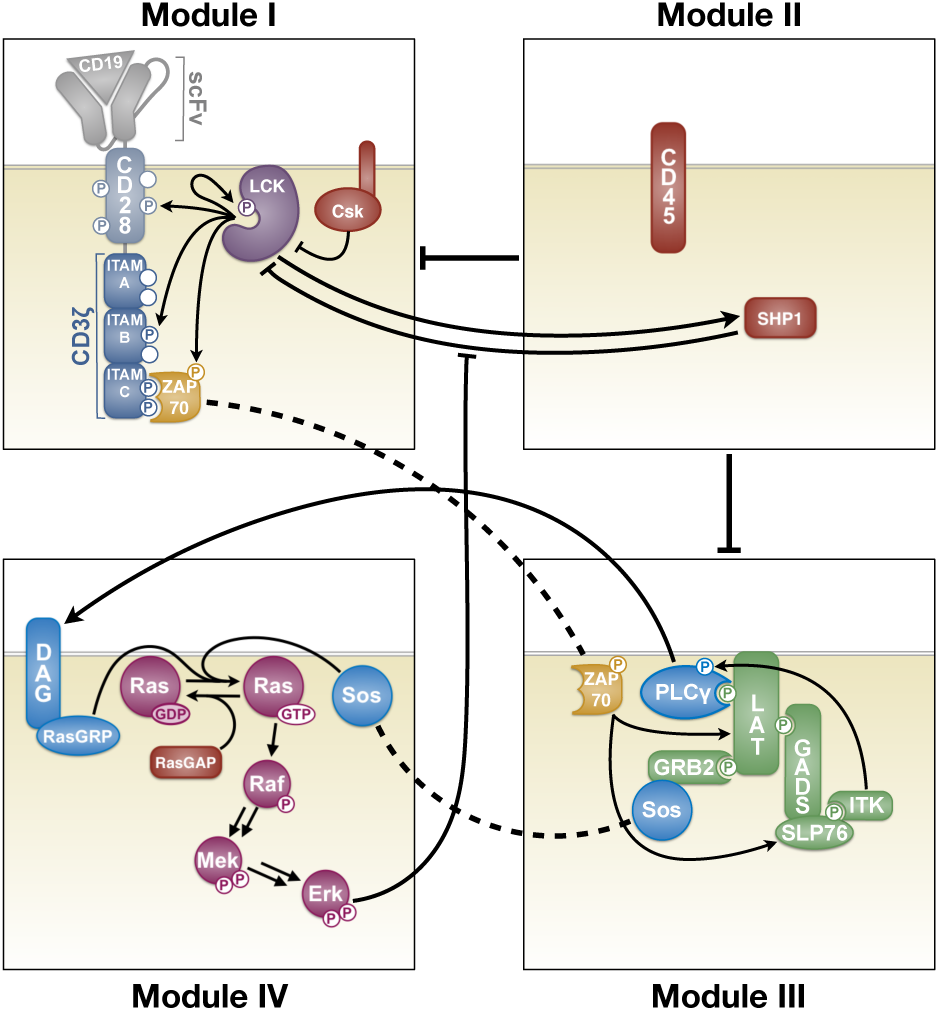
Schematic of the mechanistic model. Module I: LCK regulation, autophosphorylation, and phosphorylation. Module II: Phosphatase activity of CD45 and SHP1. Module III: Formation of the LAT signalosome and its downstream signaling. Module IV: MAPK signaling and ERK-mediated negative feedback. Arrow and bars indicate activating and inhibitory interactions, respectively. Dashed lines denote the same species in multiple Modules. Reprinted with permission from Rohrs et al. (Rohrs et al., 2019)

The model, consisting of 245 total species, 23 of which have non-zero initial conditions, and 159 parameters, was constructed using BioNetGen (Harris et al., 2016). BioNetGen is a rule-based approach for model construction that produces the set of nonlinear, coupled ordinary differential equations (ODEs) to describe how the molecular species’ concentrations evolve over time. The parameters of the model were previously fit to quantitative membrane reconstitution experiments (Hui and Vale, 2014; Hui et al., 2017; Rohrs et al., 2018) and validated using *in vitro* cellular experiments (Rohrs et al., 2019). The initial protein concentrations were taken from previous literature (Rohrs et al., 2019). Here, we simulated the model using MATLAB (MathWorks, Inc.).

### Simulating Cell-to-Cell Heterogeneity

In order to explore how heterogeneity in protein expression impacts CAR T cell activation, we performed Monte Carlo simulations to create a population of 100,000 cells. This population size was deemed large enough for the data-driven analysis described in the following sections to infer the relationships between model inputs and outputs. For each of the simulated cells, the initial protein concentrations were sampled from a log-uniform distribution over a range of 10-fold above and below the baseline protein expression used in the original model. Such a range was chosen as to encompass high and low values of protein expression to determine how cells would behave at more extreme values away from the mean. Each cell was then simulated (i.e., the model was run with each combination of the initial protein concentrations), given the same amount of antigen. The antigen concentration was set to be high enough so that it would always saturate the amount of receptor. The reason for this is that here, we examine the intracellular components of the network. Varying the antigen concentration would inevitably affect the model output, which is not the focus of the present analysis.

The model was simulated for a duration of 15 minutes. This duration was chosen because we are interested in factors affecting a rapid response to stimulation. We repeated the simulations and subsequent analysis for longer durations, 30 and 60 minutes, to see if the influential proteins change with longer stimulation.

### Characterizing Cellular Response

We used the concentration of double phosphorylated ERK (ppERK) as a way to characterize the cells’ response to antigen stimulation. Once the time course for ppERK was simulated for each cell, the final value was compared to the initial total amount of ERK in the cell. If the relative amount of ppERK reached 50% or greater, the cell was considered to have responded to stimulation and was placed into a group called “high ERK response.” Cells that failed to reach 50% of ERK phosphorylation were considered to be “low ERK response.” This threshold was chosen based on its usage in previous experimental studies (Altan-Bonnet and Germain, 2005). This classification is the primary output of the model.

### Partial Least-Squares Analysis

In brief, partial least-squares (PLS) allows for the formation of a predictive model of a system’s outputs given any number of inputs. However, unlike a mechanistic model, which describes a system based on its biological interactions, the parameters found by a PLS model do not correspond to actual biological functions. Rather, the parameters are chosen based on their ability to relate the inputs to the outputs. A PLS model can be used generalize the relationship between the inputs and the outputs, providing valuable multivariate information that would be more difficult to get out of a mechanistic model.

A PLS model is a type of multivariate regression that relates input variables to output variables by maximizing the correlation between the variables. In PLS, both the inputs and the outputs are transformed into a new dimensional space comprised of principal components (linear combinations of the inputs), and a linear regression is performed between these new variables before transforming them back to the original dimensions. When transforming the inputs into the principal components space, the dimensionality of the inputs is reduced. This allows PLS to handle noisy data and collinear inputs. Additionally, this analysis calculates how much each input contributes to the transformed variables, indicated by the weight of each input. The weights can then be examined to learn general relationships between the inputs and the outputs, and determine which inputs are most influential. For these reasons, PLS is an attractive tool for multivariate analysis of large networks (Cosgrove et al., 2010; Loiben et al., 2017; Wold et al., 2001; Wu et al., 2008).

In this study, the initial protein concentrations were used as inputs to predict which group, “high ERK response” or “low ERK response”, the cell would belong to. For this analysis, the non-linear iterative partial least-squares (NIPALS) algorithm (Geladi and Kowalski, 1986) was used for fitting the PLS model. For a detailed description of the process, see Geladi and Kowalski (Geladi and Kowalski, 1986). The inputs were taken as the log-value of the initial concentrations of the 23 proteins that have a non-zero starting value. Prior to performing the analysis, these input values were scaled by subtracting the mean of the training set and then dividing by its standard deviation. The result of this is that each input had a mean of zero and a standard deviation of one. Such scaling is important so as to eliminate the effects of highly varying input ranges on the model.

The model was trained on simulations from two-thirds of the cells, with the remaining one-third left for validating the PLS model. To determine the robustness of the PLS model, training and validation was performed 100 times, randomly shuffling the data each time before splitting into the testing and validation sets. This was performed for each possible number of principal components that the model could have. The final number of principal components was chosen as the lowest number of components where the addition of more components failed to improve the accuracy of the model.

### Identification of Influential Proteins

The primary way of identifying the most influential inputs to a PLS model is by calculating the variable importance of projection (VIP) scores. VIP scores are also used in variable selection in order to determine which variables to keep for model reduction when dealing with large numbers of inputs. VIP scores are calculated using the weights from the inputs to each component, along with the amount of output variance explained by each component. A higher VIP score indicates that the input is more influential to the outputs. Traditionally, an input is considered to be highly influential and chosen during variable selection if its VIP score is greater than one (Akarachantachote et al., 2014).

To further determine how each protein’s expression influences the response, the components of the PLS model and their relations to each protein were examined. First, looking at the components of the PLS model, we can see if high or low values of the component are associated with a particular group (“high ERK response” or “low ERK response”). Second, the absolute value of the weight for each input that makes up the component indicates how much influence that input has on a group. Finally, the sign of the weight indicates in which direction each input influences the value of the component. As an example, if high values of a component correspond to the “high ERK response” group, then inputs (initial protein concentrations) with positive weights in that component are positively associated with that “high ERK response” group. This means that increasing the values of those proteins’ initial concentrations will increase the number of cells with high ppERK levels. This examination of the PLS components and the weights of the inputs that make up the components provides a straightforward method for determining how protein expressions influence the system at the population level.

## 3 RESULTS

### 3.1 Mechanistic Model Predicts Heterogeneous Response in ERK Activation

The simulations for the population of 100,000 cells with the mechanistic model show that there is a large range of responses within the population. This is not unexpected, as the signaling responses of cells directly depend on the initial protein levels, which we explicitly varied. Both the time at which ERK activation occurs and amount of activation vary widely. Figure 2 shows the time courses for 100 randomly selected cells as a representation of the population. Approximately half of the cells in the population (42.3%) reached a level of ERK activation high enough to be classified as “high ERK response”. While there were some cells that achieved intermediate levels of ERK activation, most of the cells were at the extremes, with either almost complete activation or almost no activation. In total, approximately 91% of the cells experienced ERK activation in which ppERK was either greater than 90% or less than 10% of the cell’s initial amount of ERK. The distributions of the final relative ppERK values are shown in Figure S1A. From this, the clear “all-or-nothing” phosphorylation of ERK that is characteristic of T cells can be seen (Altan-Bonnet and Germain, 2005; Birtwistle et al., 2012). Figure S1B shows the initial concentrations of ERK that lead to “high ERK response” and “low ERK response.” From this, it is clear that the initial expression of ERK has no influence on its phosphorylation.

**Figure 2:**
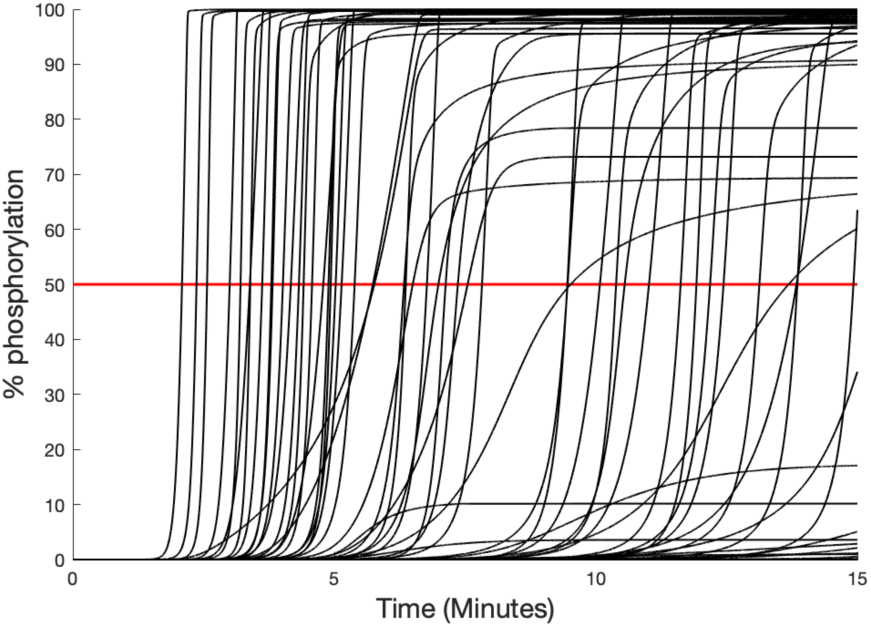
Relative ppERK time courses for a representative set of 100 cells. The red line at 50% shows the level of phosphorylation needed to be considered as a “high ERK response” following stimulation.

### 3.2 PLS Model Identifies Influential Proteins

A PLS model was developed to determine how variations in protein expression influence the ability of a CAR T cell to respond to stimulation. The inputs to the PLS model were the initial protein concentrations for each cell (see Methods), and the output was the cell’s classification as a “high ERK response” or “low ERK response.” We considered different PLS models where the number of components ranged from two to 23, which was the maximum number of components possible (the total number of inputs). The final model consisted of two components, as adding more components failed to improve accuracy in predicting the classification of the cells. Using 100 randomized sets of training and validation data, the model was able to achieve an average accuracy of 86.9% in predicting which group a simulated cell with particular initial concentrations would belong to. Considering “high ERK response” to be positive, the true positive prediction accuracy was 85.1%. The true negative prediction accuracy was 88.2%. These values are close to each other and to the overall prediction accuracy, meaning that the PLS model is not biased towards either group. This is high accuracy, considering that PLS is a linear method, while the mechanistic model that was used to produce the data is very complex and inevitably has many nonlinearities. To see how the number of training samples affected the ability of the PLS model to fit the mechanistic model, we trained the model on a smaller set of samples. Even using only 1% of the simulated population as the training set yielded similar accuracy. However, with fewer training samples, the VIP scores were less consistent between randomized batches, depending on which samples were used for training. Because of this, we moved forward with a large training set for determining the VIP scores. Overall, we established a PLS model that is able to predict which CAR-engineered T cells respond to antigen stimulation, given the initial concentrations of the intracellular signaling proteins.

Next, we used the predictive PLS model to evaluate the importance of each protein on ERK activation in the simulated CAR T cells. In order to determine which proteins hold the most influence over the response, the VIP score for each protein was calculated (Figure 3). Out of the 23 proteins in the network whose initial concentrations were varied, six achieved a VIP score of greater than one and were thus identified as being highly influential: LCK, CD3ζ, Ras, Raf, MEK, and SHP1. These proteins were then examined in detail to determine how they are influential to the system.

**Figure 3:**
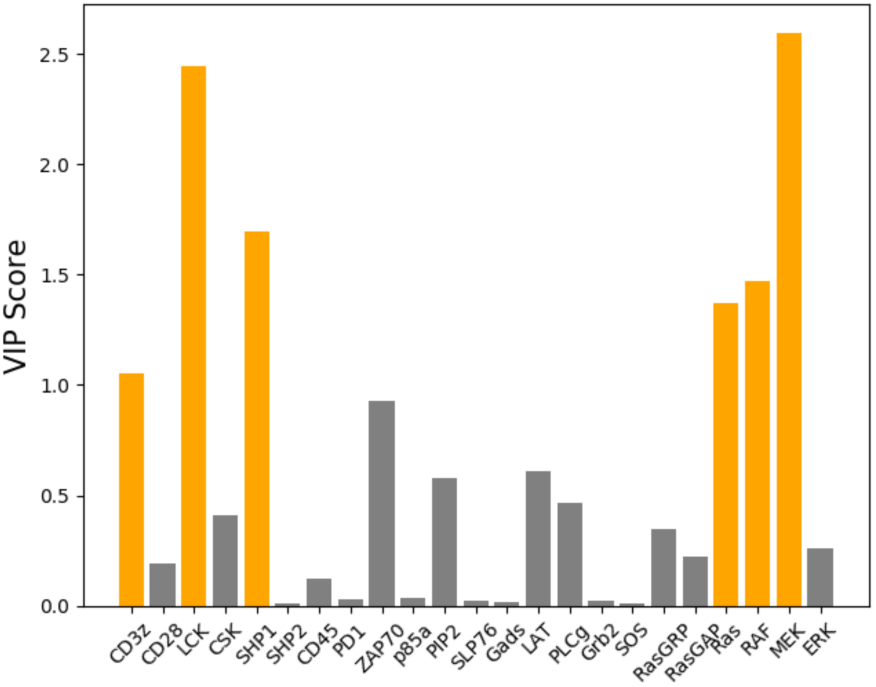
VIP scores when grouping “high ERK response” and “low ERK response”. Proteins that achieved a VIP score greater than one are colored orange, indicating that they significantly influence the PLS model’s classification of a cell being a “high ERK response” or “low ERK response”.

### 3.3 PLS Model Characterizes the Roles of the Influential Proteins

The VIP score for each input indicates whether that input significantly influences the model output. However, it does not tell whether varying the input increases or decreases the output. In order to determine how each of the six proteins with a VIP score greater than one influenced activation, the values of transformed inputs for the full model were examined. Since the optimal PLS model consisted of two components, and thus each original set of inputs was condensed into two variables, the values of the transformed inputs were viewed as a scatter plot in two dimensions (Figure 4). Each transformed input was colored based on whether it corresponded to a cell with high or low ERK response. This plot allowed for determining visually how the value of each component corresponded to the response of the cell. As shown, while there is some overlap between the two groups, there is a clear separation of “high ERK response” and “low ERK response” when considering the first component. The second component, however, provides little additional information for classifying the cells. From the PLS model, we found that the first component captures more than half (53.3%) of the variance in the output, while the second component captures very little of the variance (0.01%). Therefore, we only used the first component to determine which groups the inputs relate to. High values for the first component were associated with “low ERK response” while low values were associated with “high ERK response”. Therefore, if a protein had a positive weight for transformation into the first component, increased expression of that protein would make the cell less likely to have high levels of ppERK. The opposite is true for proteins that had negative weights. If the value of a weight was large in magnitude, then changes in the expression of its corresponding protein would have a larger effect compared to a protein whose weight for the first component is smaller in magnitude.

**Figure 4:**
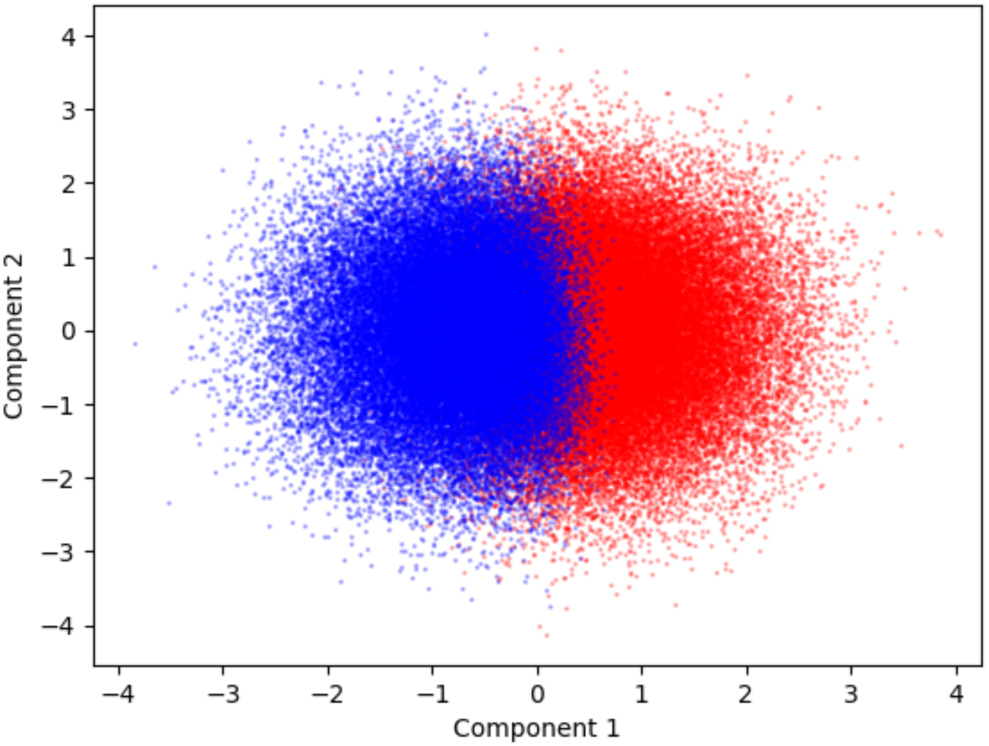
Transformed input values for each PLS model component. Blue points correspond to cells classified as “high ERK response” based on the model inputs. Red points correspond cells classified as “low ERK response” based on the model inputs.

With this insight, the weights of the first component (Figure 5) show how each of the influential proteins affects cell activation. The analysis indicates that increasing the initial amount of LCK, Ras, Raf, or MEK positively influences the cell and promotes phosphorylation of ERK. Conversely, increasing CD3ζ or SHP1 negatively influences the system, inhibiting ERK phosphorylation. We further examined the network to understand where these points of control are placed in the signaling pathway. CD3ζ, LCK, and SHP1 can all be grouped as proteins involved in the initiation of the signal, while Ras, Raf, and MEK, which make up the MAPK pathway, are all involved in conversion of the signal to a digital output. It must be noted that together, these proteins are positioned at the beginning and end of the signaling pathway.

To determine the biological reason for the influence of these six proteins, we looked into the roles that have been established in the literature. Upon binding to the antigen, specific tyrosine residues in the immunoreceptor tyrosine-based activation motifs (ITAMs) on the CD3ζ chain, which is part of the CAR, are phosphorylated by LCK. These phosphorylated ITAMs can then proceed to activate downstream signaling proteins, initiating signal transduction via ZAP-70 (Simeoni, 2017). Logically, it makes sense that increasing levels of LCK increases the cell’s ability to induce intracellular signaling, since more phosphorylation of CD3ζ would lead to a stronger signal transduction. On the other hand, upon receptor stimulation, SHP1 can bind to CD3ζ, become activated by LCK, and then proceed to dephosphorylate CD3ζ, LCK, and ZAP-70. This provides a form of negative feedback to prevent noise from accidentally leading to ERK activation (Altan-Bonnet and Germain, 2005). Therefore, increasing levels of SHP1 increases the strength of the negative feedback and prevents the signal from the receptor from being transmitted downstream. Finally, our analysis reveals that increasing CD3ζ, the last protein involved in signal initiation, inhibits cell activation. This result is discussed in detail in the following section as the conclusion drawn from the PLS model is particularly interesting and unexpected.

**Figure 5:**
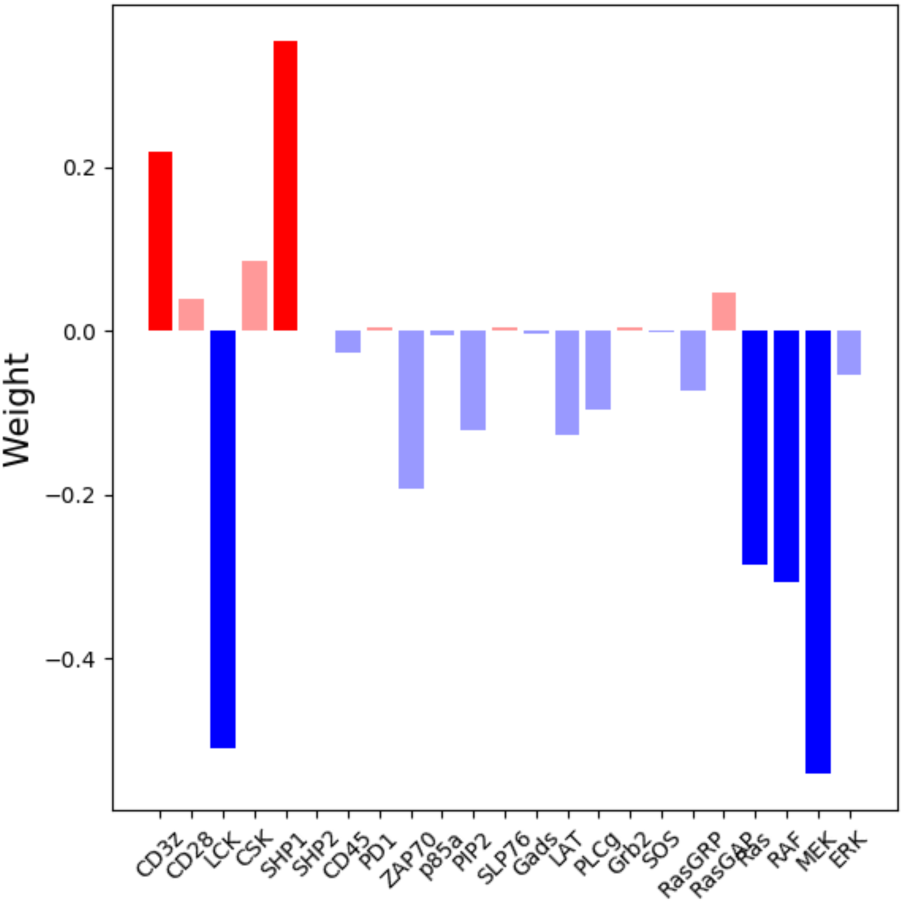
Weights from the inputs to the first component of the PLS model. Blue bars represent proteins that lower the value of the first component, thus influencing the cell to have “high ERK response”. Red bars represent proteins that increase the value of the first component, thus influencing the cell to have “low ERK response”. The proteins found to be influential have darker bars.

Ras, Raf, and MEK are three proteins at the end of the CAR-mediated signaling network and comprise the MAPK pathway, and their activation is important for the digital interpretation of a signal (Das et al., 2009). MAPK signaling leads to the rapid phosphorylation of ERK in an all-or-nothing fashion (Birtwistle et al., 2012). These actions allow the cell to make important decisions based on signals from the external environment (Shaul and Seger, 2007). As expected, analysis of the weights in the PLS model show that having increased amounts of the proteins that make up this pathway leads to more phosphorylation of ERK.

### 3.4 High Levels of CD3ζ Can Increase the ERK Response Time

Of interest is the fact that the PLS model found that increasing the expression of CD3ζ would negatively influence the cell, causing it to not respond to stimulation. At first, this seems counterintuitive, since CD3ζ binds to the antigen, becomes phosphorylated, and initiates downstream signaling. Logically, having more CD3ζ should lead to a greater ability to initiate signaling. To determine the cause for this result, we performed a series of simulations with the mechanistic model in which the initial concentrations of all of the proteins were set to their average (baseline) values, except for CD3ζ, which was varied 10-fold above and below its mean value. These simulations showed that the time of ERK activation was delayed as CD3ζ concentration increased (Figure 6A). Upon studying the predicted time courses for other proteins, we found that activated SHP1 reached higher levels when CD3ζ concentration was increased (Figure 6B). Examining the interactions involving CD3ζ and SHP1 provided an explanation for these results. Inactive SHP1 binds to CD3ζ, where it becomes activated by LCK and is then able to inactivate other molecules in the pathway. Thus, increased levels of CD3ζ allow SHP1 to become activated faster, which then inhibits downstream signaling. This analysis reveals that together, the PLS model and the details of the signaling network produce relevant insight into the cells’ response.

**Figure 6:**
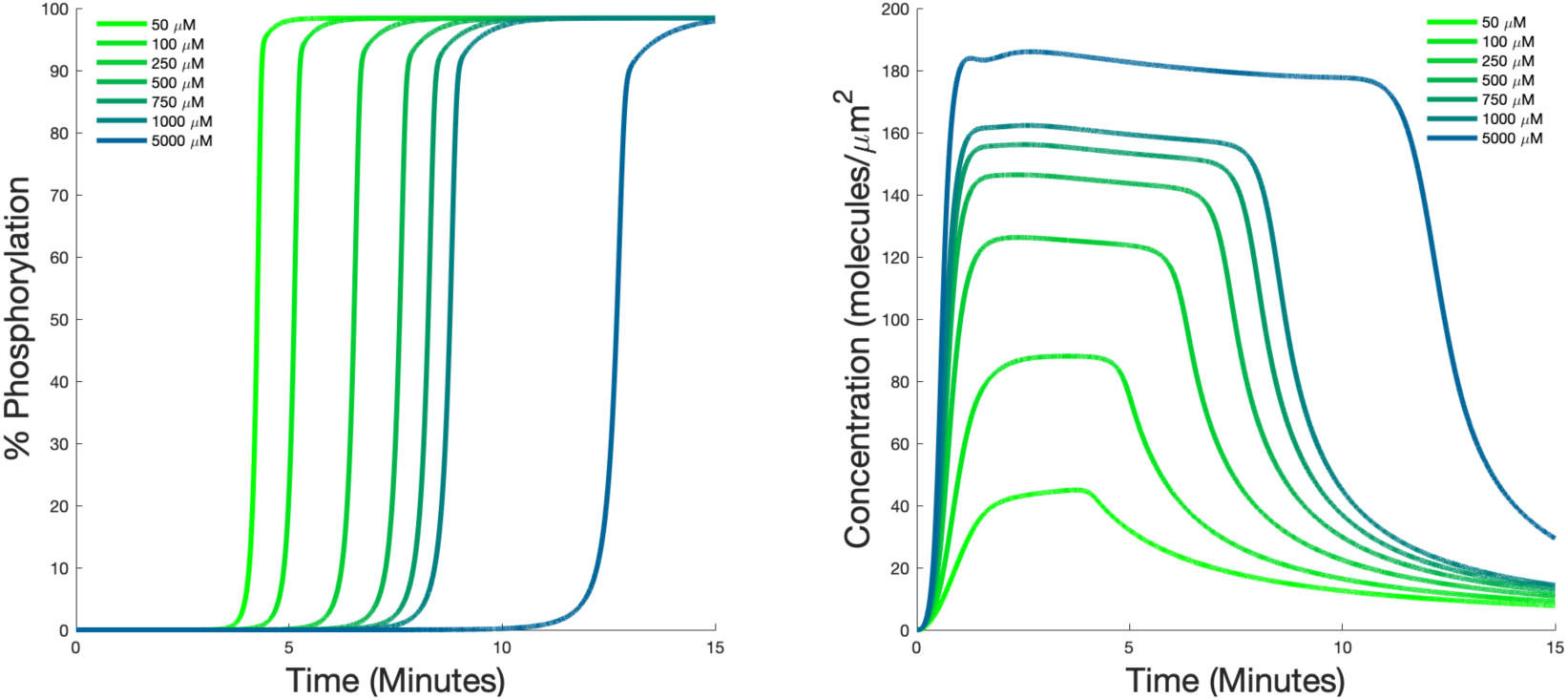
Predicted dynamics of signaling species. Time courses for phosphorylated ERK (A) and SHP1 (B) for different initial values of CD3ζ.

### 3.5 Fewer Proteins Are Influential as Simulation Time Increases

Finally, we aimed to determine whether the influential proteins change when the system is simulated for longer times. We repeated our analysis after simulating the system for 30 minutes and 60 minutes. We found that as simulation duration increased, the number of influential proteins decreased. At 30 minutes, CD3ζ ceased being influential, while the other five proteins retained their influence. At 60 minutes, LCK, RAF, and MEK were the only proteins found to still be influential. These results indicate that the key modulators of the ppERK level vary with time.

## 4 DISCUSSION

In this work, we applied partial least-squares to analyze predictions from a detailed mechanistic model of CAR T cell signaling. In particular, we study how heterogeneity in protein expression affects cell behavior characterized by MAPK signaling and phosphorylation of ERK. Through this analysis, we determined the influence of the levels of individual proteins on the ability of CAR T cells to respond to stimulation. Although the network that mediates signal transduction is fairly large and complex, we found that the expressions of only a handful of proteins play an influential role in the response. The influential proteins are positioned at the beginning and the end of the signaling pathway. The analysis was able to determine how each protein influenced the response, and while the effects of most of these proteins made sense based on their known functions, the influence of CD3ζ was found to go against what might be intuitively assumed.

Previous modeling work has also explored the role of heterogeneous protein expression on influencing cellular signaling. A study by Birtwistle *et al.* examined the response of the MAPK cascade (Birtwistle et al., 2012). They compared the result of stochastic simulation using the Gillespie algorithm to the effect of randomly sampling initial protein concentrations and found that the latter matched their experimental measurements much better than the former. This provides support to our analysis presented here. Our work expands on the general conclusion that initial protein concentrations affect cellular response and uses analysis of the mechanistic model to identify which proteins cause the observed heterogeneity. As such, our computational analysis provides novel insight that would be much more difficult to obtain experimentally.

An earlier study by Feinerman *et al.* also examined heterogeneity in T cell activation using flow cytometry to explore the influence of the expressions of CD8, ERK, and SHP1 (Feinerman et al., 2008). Similar to the results we present here, they found that increasing the initial ERK has little effect on the ability of the population to become activated, while increasing expression of SHP1 lowered the percent of the population that could respond. With our analysis, we were also able to identify additional proteins in the system that influence the response.

By gaining an understanding of which proteins in the network contribute heavily to the response to an input we can determine how an intracellular signaling network can be modulated to induce a desired response. Ideally, one would be able to modulate each cell individually to achieve optimal response; however, that is not likely feasible. The analysis done here is useful as the VIP scores, which determine how influential a protein is, and the weights of the inputs, which determine the direction in which an input influences the output, are calculated based on all of the sets of inputs. This means that the PLS model tells how the population in general would respond to an increase or decrease of a specific protein and provides information on which proteins could be modulated at the population level to attain a desired response. Moreover, by performing the data-driven analysis for different durations in the mechanistic model, it is possible identify time-based strategies for altering the cells’ responses.

There are a few limitations to using PLS for the analysis done here. The primary issue is that PLS assumes linear relationships between the inputs and outputs. It is unlikely that a system of this size is perfectly linear. However, as shown by the high accuracy of the PLS model we developed, linear relationships can be a reasonable approximation. The use of nonlinear methods, such as neural networks, would provide a higher prediction accuracy, but at the cost of being more difficult to analyze. Another limitation is that all of the initial protein concentrations were sampled from the same range. Due to the variety of different methods of gene regulation, some proteins could have wider or narrower distributions. It is possible that changing the distribution could influence the predicted numbers of each cell phenotype (i.e., high or low ERK response). However, we do not believe that this significantly impacts our analysis as we are interested in determining the effects of each protein even at higher or lower levels compared to the mean value. As more quantitative measurements for the single-cell concentrations and distributions of proteins become available, we can incorporate that information into our model.

Encoding the outputs of a mechanistic model as a data-driven model, which reduces the computational time, has multiple uses. As described in this work, the data-driven model can then be used as an analysis tool. Here, the data-driven model enabled a better understanding of the important relationships between model variables and cellular response. Those relationships can inform how to engineer the cell for a desired purpose. Another potential application is using the data-driven model inside of an agent-based model (ABM). While some ABMs do use simple ODE models as a way of making cellular decisions (Hendrata and Sudiono, 2016; Wang et al., 2007; Zhang et al., 2009), most use discrete or probabilistic rules to govern how each cell behaves, as that is much more computationally efficient. Using a data-driven model, ODE networks could potentially be simplified, allowing ABMs to become more biologically detailed without a significant increase in computational cost.

The results from this study show that data-driven analyses provide insight into large mechanistic models. Using a relatively fast analysis, we were able to determine which proteins were the most influential in determining the response of the system, and whether each protein had a positive or a negative influence. Using PLS provides information on how to push the population as a whole towards a specific response. In the context of CAR T cell signaling, we found that the system was most sensitive to proteins at the very beginning or very end of the network. While most of the proteins (LCK, SHP1, Ras, RAF, and MEK) influenced the system in a way that is expected based on their biological functions, we found that CD3ζ can actually influence the system towards no response, despite being part of the receptor that initiates signaling. This result was explained by examining the specific interactions in the mechanistic model. Overall, we find that data-driven methods are capable of analyzing detailed signaling networks, rather than just being used in cases where forming a mechanistic model is not feasible.

## 5 CONLCUSION

Here we used a data-driven analysis of a mechanistic model to study how variations in protein expression influence the ability of a CAR T cell to respond to stimulation and promote ERK phosphorylation. We identified six proteins relating to either the receptor or the MAPK cascade that strongly influenced the output of the system. We also found the counterintuitive result that increasing the amount of receptor in the system can actually hinder ERK phosphorylation, as it increases the level of active phosphatase in the system. By combining data-driven and mechanistic modeling, we gain useful insight into cell signaling.

## ACKNOWLEDGEMENTS

The authors thank Lauren Slowskei for initial work on this project and the members of the Finley research group for their critical feedback. This work was supported by The Graduate School Provost Fellowship (to CGC).

## SUPPORTING INFORMATION

**Figure S1.** Histograms of final relative ppERK concentrations and initial ERK concentrations.

## AUTHOR CONTRIBUTIONS

SDF conceived of the work. CGC completed the simulations and analysis. Both authors wrote and edited the manuscript.

