## Supplementary material for "Data-Driven Analysis of a Mechanistic Model of CAR T Cell Signaling Predicts Effects of Cell-to-cell Heterogeneity": Figure S1

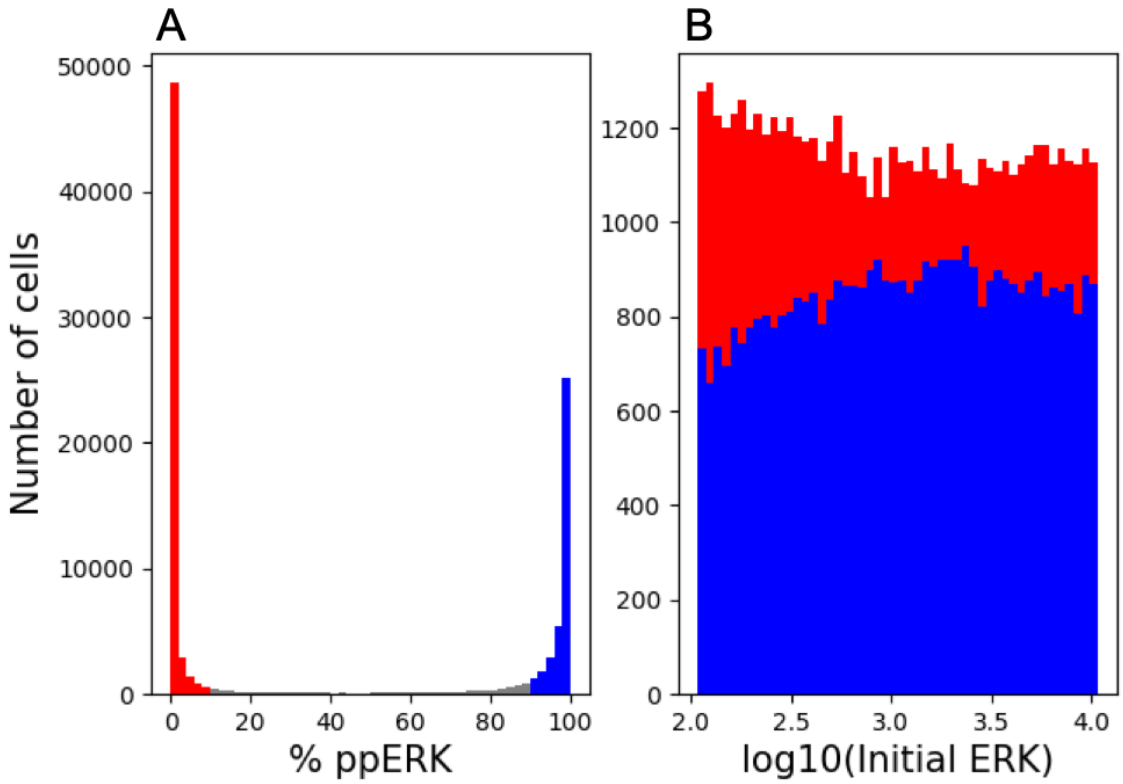


**Figure S1:** (A) Histogram of the final relative concentrations of phosphorylated ERK. Concentrations below 10% are shown in red, above 90% in blue, and in between in gray. (B) Histogram showing distributions of initial ERK concentrations that led to a response (blue) and no response (red).
